# Genome-wide evolutionary response of European oaks since the Little Ice Age

**DOI:** 10.1101/2021.05.25.445558

**Authors:** Dounia Saleh, Jun Chen, Jean-Charles Leplé, Thibault Leroy, Laura Truffaut, Benjamin Dencausse, Céline Lalanne, Karine Labadie, Isabelle Lesur, Didier Bert, Frédéric Lagane, François Morneau, Jean-Marc Aury, Christophe Plomion, Martin Lascoux, Antoine Kremer

**Affiliations:** UMR BIOGECO, INRAE, Université de Bordeaux, 69 Route d’Arcachon, 33612 Cestas, France; College of Life Sciences, Zhejiang University, Hangzhou, Zhejiang 310058, China; Department of Botany & Biodiversity Research, University of Vienna, Vienna, Austria; Genoscope, Institut de Biologie François Jacob, Commissariat à l’énergie atomique (CEA), Université de Paris-Saclay, 2 rue Gaston Crémieux, 91057 Evry, France; Helix Venture, 33 700 Merignac France; Département Recherche Développement Innovation, Office National des Forêts, 100 Boulevard de la Salle, 45760 Boigny-Sur-Bionne, France; Génomique Métabolique, Genoscope, Institut François Jacob, CEA, CNRS, Univ Evry, Université Paris-Saclay, 91057 Evry, France; Department of Ecology and Genetics, Evolutionary Biology Centre, Uppsala University, 75236 Uppsala, Sweden

**Author notes:** These two authors contributed equally to the work (Dounia Saleh, Jun Chen). Service de l’Information Statistique Forestière et Environnementale, Institut National de l’Information géographique et Forestière, Château des Barres, 45290 Nogent sur Vernisson France (François Morneau).

**Keywords:** Linked selection, evolution, *Quercus petraea*, Little Ice Age

## Abstract

The pace of tree microevolution during Anthropocene warming is largely unknown. We used a retrospective approach to monitor genomic changes in oak trees since the Little Ice Age (LIA). Allelic frequency changes were assessed from whole-genome pooled sequences for four age-structured cohorts of sessile oak (*Quercus petraea*) dating back to 1680, in each of three different oak forests in France. The genetic covariances of allelic frequency changes increased between successive time periods, highlighting genome-wide effects of linked selection. We found imprints of convergent linked selection in the three forests during the late LIA, and a shift of selection during more recent time periods. The changes in allelic covariances within and between forests mirrored the documented changes in the occurrence of extreme events (droughts and frosts) over the last three hundred years. The genomic regions with the highest covariances were enriched in genes involved in plant responses to pathogens and abiotic stresses (temperature and drought). These responses are consistent with the reported sequence of frost (or drought) and disease damage ultimately leading to the oak dieback after extreme events. Our results therefore provide evidence of selection operating on long-lived species during recent climatic changes.

## INTRODUCTION

Our ability to forecast the response of species to climate change is limited by our lack of knowledge about the pace of adaptive evolution, particularly in long-lived species, such as trees. Standing tree populations already had to survive important environmental changes during their lifetime and these environmental challenges likely triggered significant evolutionary changes, though the magnitude of tree adaptive response remains to be quantified. More specifically, multicentennial trees in the Northern Hemisphere that are old enough to have experienced the transition from the cold Little Ice Age (LIA, 1450-1850) to the warm Anthropocene (1850-today) (Luterbacher *et al*. 2004; Corona *et al*. 2010; Luterbacher *et al*.2016; Anchukaitis *et al*. 2017) offer a unique opportunity to assess the extent of genome-wide response to this extreme and well described environmental challenge. The LIA (Tkachuck 1983) was a cold period characterized by climatic extremes such as long and harsh winters, but also severe droughts that led to plagues, famines and ultimately revolutions (Pfister 1984; Fagan 2002; Parker 2013). The consequences of the LIA for plants are best illustrated by the recurrent poor crop harvests (Le Roy Ladurie 2004, 2006). Evidence of the impact of the LIA on forest trees is provided by comparisons of tree ring sizes and historical temperature records (Carrer & Urbinati 2006; Edouard *et al*. 2009). According to inferences concerning past forest tree dynamics from subfossilized wood remains, the decrease in temperature during the LIA resulted in a retreat of the forest tree line at high latitudes (Kullman 2005; MacDonald *et al*. 2008; Linderholm *et al*. 2014; Kullman 2015; Helama *et al*. 2020) or altitudes (Camarero *et al*. 2015) and changes in species composition (Campbell & McAndrews 1993). Here, we investigated whether climatic trends during and after the LIA and occurrences of extreme events had evolutionary consequences for forest tree populations. We addressed two major questions in this work. First, we investigated whether extreme events occurring during the late LIA left a genomic signature. Second, we investigated whether the shift in climate after the LIA also left a genomic imprint. We chose sessile oak (*Quercus petraea* (Matt.) Liebl.) as the model species for this study, as the oldest forests in Europe contain sessile oak stands that came into existence in the middle of the LIA (near 1650), a time at which the French statesman Colbert implemented even-aged management in French forests (Gallon 1752). Four age-structured cohorts of roughly 340, 170, 60 and 12 years old were sampled within three forests, to explore changes in allele frequencies over time. *Quercus petraea* is known to display considerable genetic diversity (Mariette *et al*. 2002; Kremer & Hipp 2020; Leroy *et al*. 2020) and to have a high genetic variance for fitness (Alexandre *et al*. 2020). The selection for viability or the demographic dynamics generated by extreme weather events would be expected to result in changes in allelic frequency. Climate change-driven evolution over the course of a few generations has mostly been reported in invasive species (Chown *et al*. 2015) or in controlled experiments (Ravenscroft *et al*. 2015). By contrast, in this study, we explored the ability of a native species with high levels of standing genetic variation to respond to recent documented climatic changes. Our approach mirrors experiments monitoring the change in allelic frequencies over successive discrete generations *in natura* (Malaspinas 2016) or in controlled selection experiments (Schlotterer *et al*. 2015), except that our study made use of contemporary age-structured cohorts. Earlier genome-wide investigations performed with a synchronous approach in common gardens highlighted the multifaceted and unrepeatable signatures of natural selection, characterized by heterogeneous polygenicity (trait architecture determined by a large number of genes with small effects) and allelic heterogeneity (Alberto *et al*. 2011; Plomion *et al*. 2016; Rellstab *et al*. 2016). In two recent publications Buffalo and Coop (2019, 2020) showed that allele frequency trajectories can be shifted due to linked selection between selected and neutral loci and generate covariances between allelic frequency changes at successive time periods. Theoretically, the magnitude of covariances depends on the genetic variance of fitness, recombination and linkage disequilibrium between selected and neutral loci, whereas their sign depends on the maintenance or fluctuation of selection pressures over time. We hypothesized that this approach would be suitable for detecting genomic footprints of selection in the past, during the late LIA, and shifts in selection pressures due to warming after the end of the LIA. We also explored a more qualitative approach addressing the underlying functions of the multiple genes contributing to the covariances of allelic frequency changes. We thus had three objectives: (1) to retrace trends for the covariances of allelic frequencies between age-structured cohorts spanning the last three centuries in oak stands (2) to determine whether these trends were repeatable over replicated observations in three different forests, (3) to explore the gene networks involved in these responses to environmental change.

## METHODS

### Sampling forests and age structured cohorts

We sampled three oak forests located in the central and western part of France (Bercé, Réno-Valdieu and Tronçais, Figure 1A). These forests include stands of up to 349 years of age when the study started in 2014, and are managed under even-aged silvicultural regimes (Supporting Information S1). The upper canopy consisted principally of *Quercus petraea*, and historical records and genetic evidence (based on chloroplast DNA haplotypes) indicated that the three forests were of natural origin (Petit *et al*. 2002). In each forest, we sampled individuals belonging to four age-class cohorts corresponding to ages of 340, 170, 60 and 12 years (born approximatively in 1680, 1850, 1960 and 2008) (Figure 1, Table S1). These cohorts are referred to as cohorts 4, 3, 2 and 1, respectively, below. The regeneration period, during which mature trees mate and the stand is renewed by natural seeding, takes today about 10 to 20 years, but extended over longer periods in the past (up to 30 years). The age of the trees in a given cohort may, therefore, vary by up to 10 to 30 years. Cohorts were dated on the basis of management records, together with dendrochronological recordings for a few felled trees within each cohort (Figure 1B). Knowledge of the demographic dynamics of even-aged forests is required to identify the periods during which selection was at its strongest. When a stand is renewed by natural seeding, a very dense cohort of seedlings develops (more than 100,000 seedlings/ha), the number of plants gradually decreasing to about 4,000/ha by the age of 10 years, as a result of natural selection and chance events. Crucially, subsequent silvicultural thinning is applied to only the remaining 6% of the trees. Hence, an oak stand that is about 340 years old today probably underwent its strongest bout of natural selection in the late seventeenth century, when it was at the seedling stage. Such reasoning provided the rationale for sampling age-structured cohorts in even-aged stands for retrospective monitoring of the selective impact of past climatic changes. The different cohorts within a given forest are probably derived from the same founding population established more than 10,000 years ago (Giesecke 2016; Giesecke & Brewer 2018), but there is no direct traceable generation-to-generation link between the cohorts.

**Figure 1.**
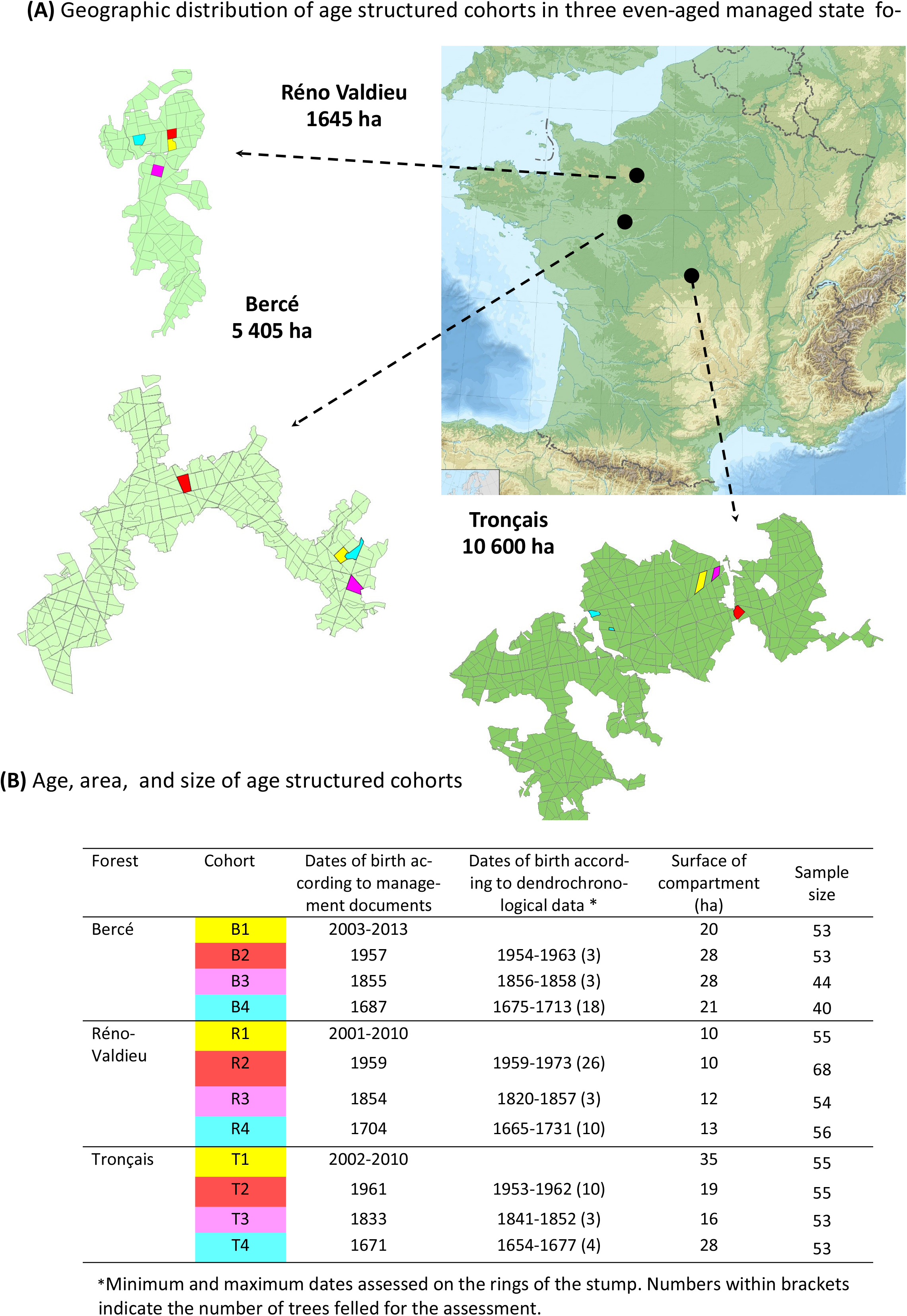
Sampling of forests and age structured cohorts of sessile oak (A) Distribution of age structured cohorts of sessile oak in three even-aged managed national forests in France. Each forest is subdivided in compartments (about 20ha in size) limited by the black lines on the forest maps. Age class compartments are evenly distributed in the forests. Densities are extremeley high at the seedling stage (>100,0000/ha) and decrease very rapidly due to natural selection during the early stage (≈ 4,000 at age 10). (B) Age, area, size of age structured cohorts. Dendrochronological data of tree rings on felled trees in each cohort allowed to confirm documentary records of trees ages. About 50 trees were randomly sampled in each cohort for whole genome sequencing.

High-resolution regional and temporal temperature reconstructions based on a combination of instrumental data (for the most recent period), documentary records, ice core and tree ring proxy data revealed clear temperature trends over the last 340 years in the three forests (Luterbacher *et al*. 2004; Luterbacher *et al*. 2016) (Supporting Information S2, Figure S1). There were two cooler periods within the LIA (1660 to 1690 and 1840 to 1860), followed by a continuous increase in temperature of about 0.8°C from 1860 to 1950. The two cooler periods were also dominated by more frequent extreme winters and summer drought events (Figure S2 and S3, Figure 3d). Comparing these temporal bioclimatic trends with the ages of our cohorts, the two oldest cohorts are almost synchronous with the two periods in which conditions were most severe during the LIA. Cohort 4 originated during the late 17^th^ century, whereas the trees of cohort 3 were established in the mid-19^th^ century (Figure 1B).

### DNA extraction and sequencing

Leaf samples were collected from cohort 1 in the spring and summer 2014, whereas cambium samples were harvested at the base of the trunk with a punch for the three older cohorts. The sample size for each cohort was between 40 and 68 trees (Figure 1B). DNA was extracted from the 639 samples with a Qiagen extraction kit. DNA yields were measured with a NanoDrop 1000 spectrophotometer (NanoDrop Technologies, Wilmington, DE, USA) and DNA samples were pooled in equimolar amounts within each cohort from each forest. The 12 pools (3 forests * 4 cohorts) were sequenced on an Illumina HiSeq4000 sequencer generating 150 bp paired-end reads.

### SNP discovery

The various steps in the SNP calling pipeline were as described in (Altmann *et al*. 2012; Pfeifer 2017). The adapters and primers were removed and reads were trimmed to remove nucleotides with a quality value below 20 from the two ends. The sequences between the second unknown nucleotide and the end of the read were removed, and reads of less than 30 nucleotides in length were discarded. Finally, the read pairs from the low-concentration spike-in Illumina PhiX Control library were removed. We obtained a mean of 348,863,070 reads per pool (Table 1). The reads were mapped onto the v2.3 *Q. robur* genome assembly (Plomion *et al*. 2018) with bwa-mem, with a seed size of 39 (Li & Durbin 2010). Incorrectly paired reads and reads giving multiple alignments were removed with samtools (Li *et al*. 2009). Duplications were removed with Picard tools (no publication, Broad Institute). Base Alignment Quality (BAQ) was calculated with samtools (Li *et al*. 2009). Pileup files were generated for each scaffold over all forests, with samtools. These files were converted into synchronized pileups with a minimum alignment quality of 10, and allele frequencies were calculated for SNPs with a minimum count of two for minor alleles, a minimum coverage of 40X and a maximum coverage of 10% of total coverage within each pool, with Popoolation2 (Kofler *et al*. 2011). Further filtering was applied to the three types of pileup files, to select biallelic SNPs with a minimum minor allele frequency of 0.02.

**Table 1.** SNP diversity statistics of the age structured cohorts

| Forest | Cohort | Number of reads after<br>post-processing | Number of<br>SNPs | $\pi \pm$ standard deviation |
| --- | --- | --- | --- | --- |
| Bercé | B1 | 427,365,137 | 13,277,388 | $0.01202 \pm 5.10^{-5}$ |
| | B2 | 444,637,716 | 13,533,867 | $0.01208 \pm 5.10^{-5}$ |
| | B3 | 450,468,871 | 13,586,680 | $0.01334 \pm 8.10^{-5}$ |
| | B4 | 439,841,819 | 13,334,778 | $0.01447 \pm 1.10^{-6}$ |
| Réno-Valdieu | R1 | 421,851,747 | 13,344,026 | $0.01211 \pm 5.10^{-5}$ |
| | R2 | 443,414,110 | 13,678,790 | $0.01256 \pm 5.10^{-5}$ |
| | R3 | 435,807,782 | 13,518,651 | $0.01203 \pm 5.10^{-5}$ |
| | R4 | 440,543,051 | 13,605,834 | $0.01206 \pm 5.10^{-5}$ |
| Tronçais | T1 | 543,901,014 | 15,592,854 | $0.01351 \pm 7.10^{-5}$ |
| | T2 | 440,923,574 | 13,457,837 | $0.01208 \pm 5.10^{-5}$ |
| | T3 | 437,816,360 | 13,704,258 | $0.01215 \pm 5.10^{-5}$ |
| | T4 | 432,251,274 | 13,514,365 | $0.01208 \pm 5.10^{-5}$ |

### Diversity within and between cohorts

Genetic diversity was estimated on non-overlapping genomic windows of 10kb spanning the whole oak genome. For each pool, we therefore generated new pileup files, including monomorphic sites. To take into account the variance in pool size and coverage among populations, we used the *subsample-pileup.pl* script from Popoolation2 (Kofler *et al*. 2011) to target a minimum coverage of 30 at each position and for each pool. These parameters were chosen after optimization in order to minimize the amount of missing data between pools. Tajima’s *π* (Tajima 1989) was then calculated with the *variance-sliding.pl* script. As for the SNP sets, sites with a minimum alignment quality of 10 were retained, and were considered polymorphic if at least two copies of the minor allele were detected among all reads. Windows with a minimum covered fraction of 50% were retained for the calculation of Tajima’s *π*. *F_ST_* values were calculated for each SNP, between each pair of cohorts, using Popoolation2 (Kofler *et al*. 2011), and were averaged over the whole genome.

### Temporal covariances

Buffalo and Coop (2019, 2020) have developed a method for testing for genomic signals of selection on polygenic traits due to linked selection, based on temporal covariances of allelic frequency changes. Allele frequency changes were calculated for different time spans separating the cohorts in the three replicated forests. We used the CVTK software package available from http://github.com/vsbuffalo/cvtk to calculate the temporal covariances between allelic frequency changes (Buffalo & Coop 2020).

We filtered SNPs by removing sites with a depth below the number of alleles sampled for each cohort, and with a minor allele frequency below 0.02. Contigs were also filtered out when shorter than 200 kb long (excluding ‘N’s). Read depth and sample size were used to correct for bias in variance estimates (Buffalo & Coop 2020). Corrections were also done for bias caused by sampling noise common to adjacent time points. This bias stems from the allele frequency at a given time point common to the terms of the covariances between adjacent time periods (e.g., for Cov(Δ_1680–1850_, Δ_1850–1960_), allele frequency at 1850 is included in both terms).

Genome-wide temporal covariances were then computed between pairs of non-overlapping time spans within each forest. Covariances were calculated based on all SNPs, and for all three forests separately. The mean and 95% confidence intervals of the covariance were obtained by bootstrapping, with 5000 iterations.

The change in the variance of allelic frequency changes over the whole period, from 1680 to 2008, becomes:

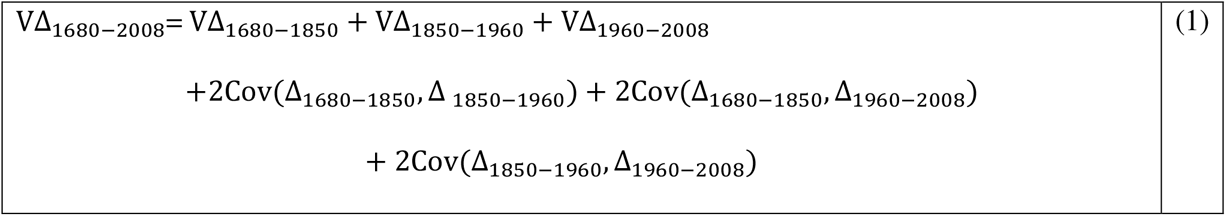

The contribution of the temporal covariances to VΔ_1680–2008_ due to linked selection thus becomes (G equation [1] of Buffalo & Coop (2020)) :

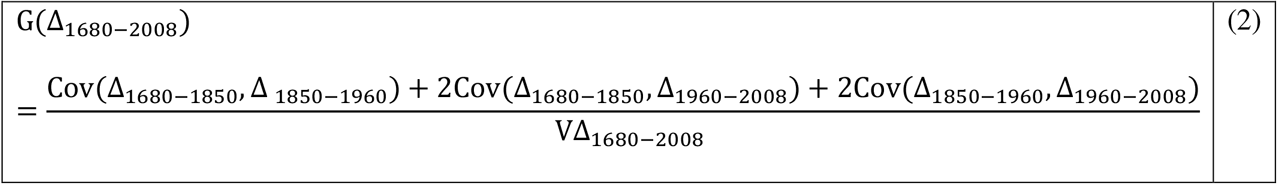

G estimates the contribution of linked selection to the variance of the allelic frequency change from the starting point to the most recent time period. Buffalo and Coop (2020) suggested that G can also be understood as the decrease in neutral diversity due to linked selection.

We extended this approach to the calculation of covariances between pairs of forests, (Cov ΔX_i-j_, ΔY_i-j_) over the same period (i-j) or at different non-overlapping time periods. The maintenance of a positive covariance would indicate similar responses and directions of selection (convergent selection) in the different forests, whereas negative values would provide evidence of differences in the direction of selection. The extension to different forests made it possible to estimate covariance over contemporary time periods, or disconnected time scales (adjacent or distant), thereby making it possible to determine whether convergent selection occurred over the same time period, or whether fluctuating selection occurred over different time periods.

### Gene annotation and ontology enrichment analysis

We divided the oak genome into tiles of 100,000 bp in length and calculated the variance and covariance for all tiles. Genomic regions with strong responses to linked selection were identified on the basis of covariance values between time intervals above 0.01 for all three replicated forests.

The genes located in the regions with the highest covariances were annotated with various bioinformatic tools. *Arabidopsis thaliana* was initially used as a reference. Orthologous genes between oak and *A. thaliana* were identified for each protein sequence (file “Qrob_PM1N_CDS_aa_20161004.fa.gz”, available at https://urgi.versailles.inra.fr/download/oak/) by selecting the best blast hit (blastP, pvalue < 1e+05) to represent the oak gene model, and *Arabidopsis* GO terms from TAIR were used for annotation (https://www.arabidopsis.org/). For oak genes without orthologs in *Arabidopsis*, we performed eggNOG-mapper v2 functional annotation based on fast orthology assignments, using precomputed eggNOG v5.0 clusters and phylogenies (Huerta-Cepas *et al*. 2017; Huerta-Cepas *et al*. 2019). Gene ontology (GO) terms were inferred from eggNOG orthologous groups (OGs).

Gene sets associated with Biological Processes (BP), Molecular Functions (MF), and Cellular Components (CC) were tested for GO terms enrichment with the “topGO” R package and the “weight” algorithm associated with a Fisher’s exact test to select the most relevant terms (Alexa *et al*. 2006; Alexa & Rahnenfuhrer 2020). A *p*-value < 0.05 was applied for the statistical test, and no FDR was calculated because the *p*-values returned by the “weight” method are interpreted by Alexa & Rahnenfuhrer (2020) as corrected or not affected by multiple testing.

## RESULTS

### Diversity and differentiation within and between age-structured cohorts

On average, more than 13 millions of SNPs were called in each cohort (Table 1). Within age-structured cohorts, nucleotide diversity was high (*π* ~0.01205). Levels of diversity were similar across cohorts and forests and consistent with previous estimates for oak (Plomion *et al*. 2018). The mean pairwise *F_ST_* between cohorts ranged from 0.010 to 0.015, with no detectable structure between cohorts and forests (Figure 2). Using Fisher exact tests, we found significantly differentiated SNPs between the consecutive cohorts 4 and 3, cohorts 3 and 2, cohorts 2 and 1 and between the distant cohorts 4 and 1 in all three forests for a limited number of SNPs in comparison to the total number of 13 millions SNPs. In Bercé the number of significantly differentiated SNPs between any two cohorts varied between 293 and 538. The range of variation was 20 to 40 in Réno-Valdieu and 45 to 1674 in Tronçais. None of these SNPs was significantly differentiated between cohorts in more than one forest. Overall these figures suggest that genome-wide differentiation between cohorts was likely due to subtle changes of allelic frequencies between different time periods.

**Figure 2.**
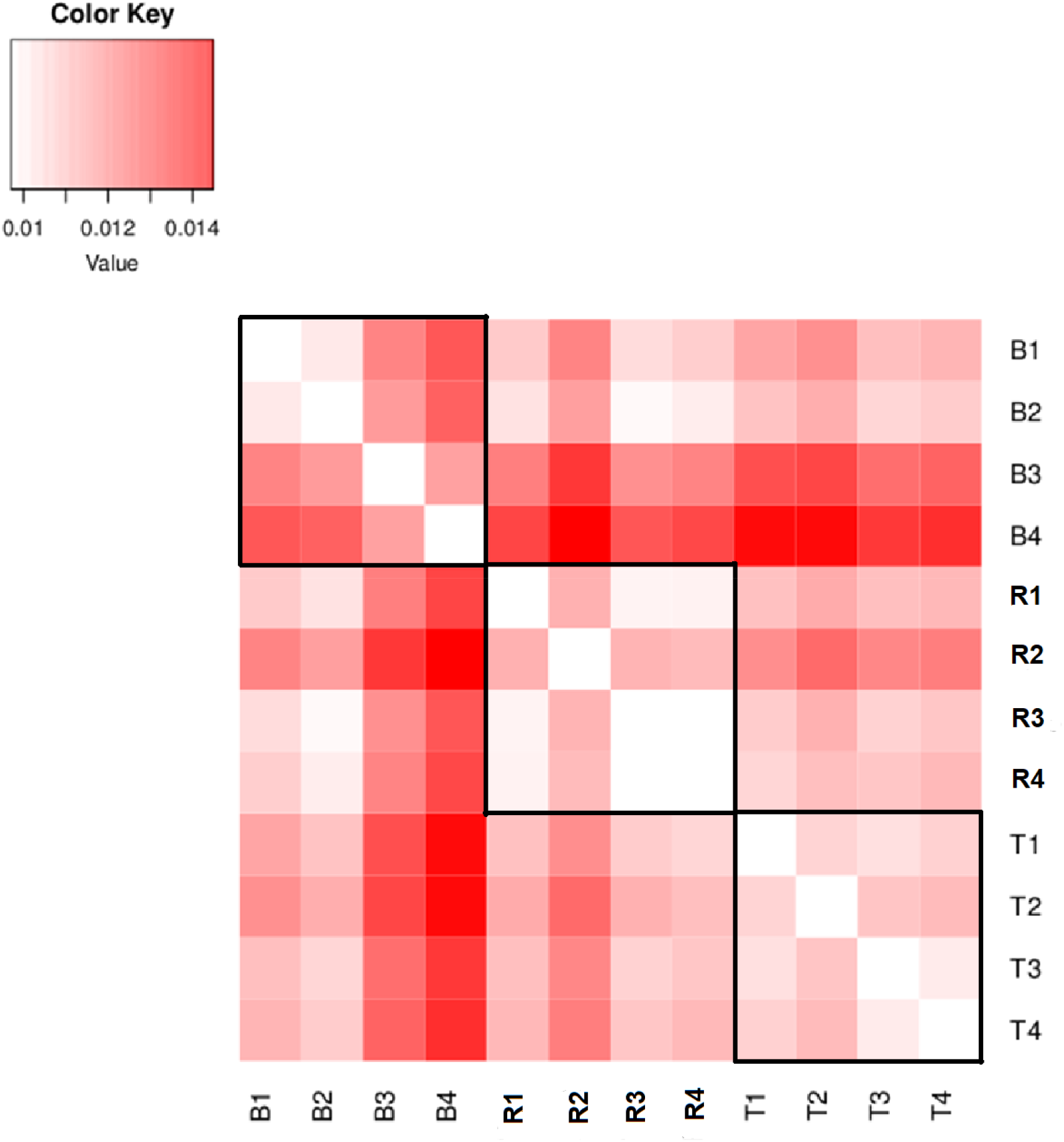
Fst values between age-structured cohorts in the three forests (B: Bercé; R: Réno-Valdieu; T: Tronçais). Subscripts to forest acronyms indicate the ages of the cohorts: 4: age ~340 or year of birth ~1680 3: age ~170 or year of birth ~1850 4: age ~60 year of birth ~1960 1: age ~12 or year of birth ~2008

### Temporal covariances of allelic frequency changes within forests

We calculated the covariances of allelic frequency changes between proximal time periods (COV(Δ_1680–1850_, Δ_1850–1960_) and Cov(Δ_1850–1960_, Δ_1960–2008_)) and distant time periods (Cov(Δ_1680–1850_, Δ_1960–2008_)). All proximal and distant covariances were significant in all three forests (Figure 3). The covariances between the earliest adjacent time periods (Cov(Δ_1680–1850_, Δ_1850–1960_) were positive and significant in each forest (Figure 3A). The covariances between the more recent adjacent time periods Cov(Δ_1850–1960_, Δ_1960–2008_) varied between the three forests, being positive in Bercé, slightly lower in Réno-Valdieu and negative in Tronçais (Figure 3B). Finally, the covariances between distant time periods were much lower than those between adjacent time periods in Bercé and Réno-Valdieu, and also partially in Tronçais (Figure 3C). Overall, the patterns illustrated in Figure 3 show a shift of covariances between adjacent time periods from the earliest periods considered (1680-1850, 1850-1960) to the more recent time periods (1850-1960, 1960-2008).

**Figure 3.**
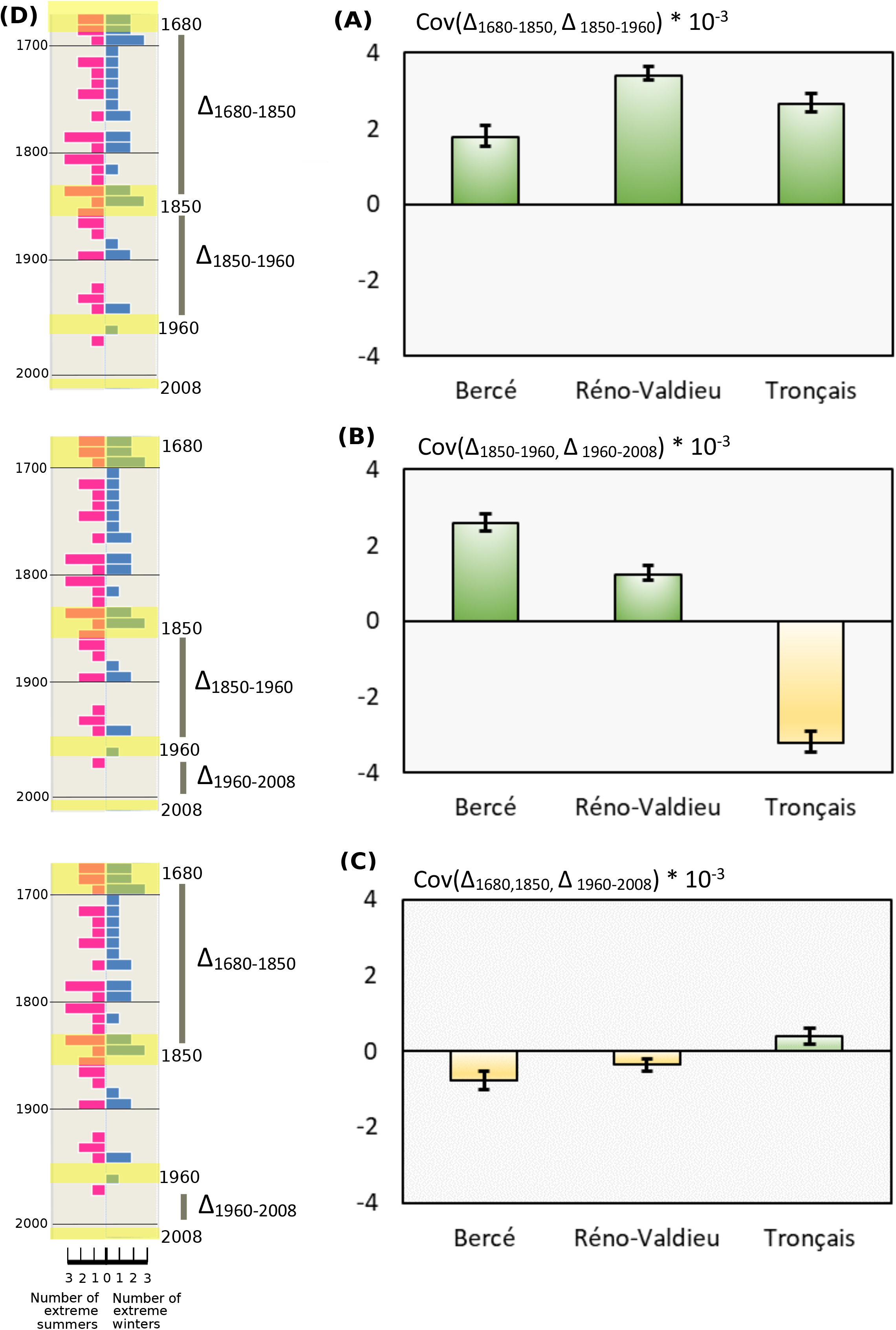
Temporal covariances of allelic frequency changes between different time periods and occurrences of extreme climatic events since the Little Ice Age. Mean and 95% confidence intervals of the covariance were obtained by bootstrapping with 5000 iterations. (A) Temporal covariances of allelic frequency changes between 1680-1850 and 1850-1960 in the three forests. (B) Temporal covariances of allelic frequency changes between 1850-1960 and 1960-2008 in the three forests. (C) Temporal covariances of allelic frequency changes between 1680-1850 and 1960-2008 in the three forests.. (D) Timeline subdivided in decades. On the right side of the timeline in blue bars, number of extreme winters per decade according to instrumental temperatures recorded at the Observatory of Paris between 1676 and 2010 (Rousseau 2012) (More details in Figure S22). On the left side of the timeline in red bars, number of extreme summer droughts per decade according to Cook’s data base of Old World megadroughts (Palmer 1965; van der Schrier *et al*. 2013; Cook *et al*. 2015) (for more details Figure S3). Highlighted decades in yellow correspond to periods when the cohorts became installed after natural regeneration.

We also estimated the contribution of the temporal covariances to VΔ_1680–2008_ by calculating GΔ_1680–2008_ (equation 2, Table 2). The proportion of the variance of allelic frequency change due to linked selection increased from time period 4 (1680) to time period 1 (2008), due to the positive covariances in Bercé (9 to 16%) and Réno Valdieu (18% to 22%). It decreased in Tronçais from 12% to −1% (Table 2), due to the overall decrease in covariances between distant time periods. The largest contribution was that of the covariances for the earliest periods Cov(Δ_1680–1850_, Δ_1850–1960_) which were positive in all three forests, resulting in GΔX_1680-1960_ values of 9%, 18% and 12%, respectively..

**Table 2.**
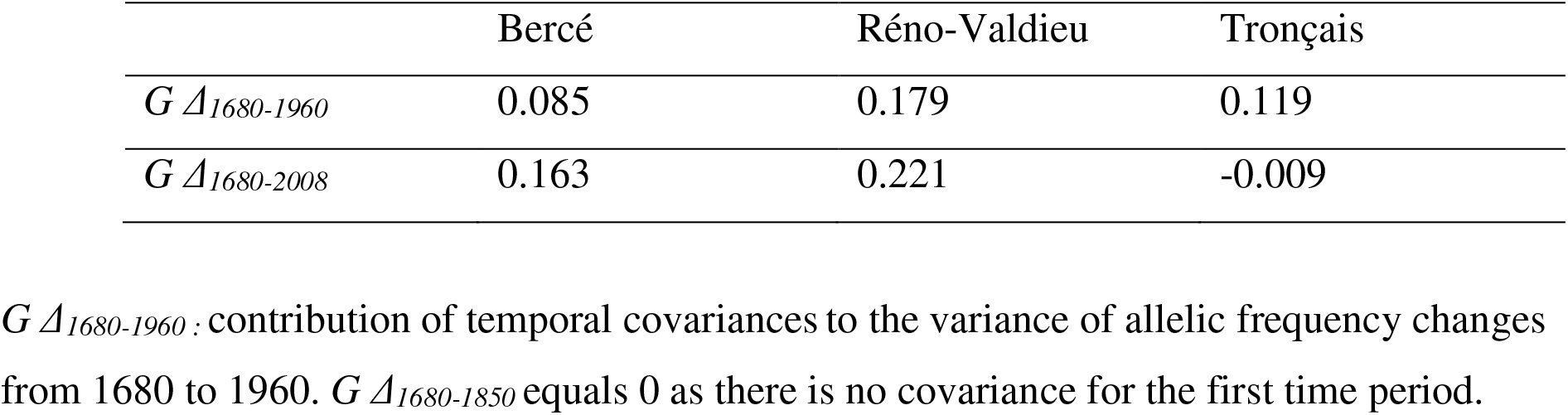
Contribution of the covariances between allelic frequency changes to the variance of allelic frequency changes between two time points.

### Temporal covariances of allelic frequency changes between forests

Significant positive temporal covariances between forests would indicate convergent linked selection. Considering different time periods for calculating the covariances can provide an indication as to when convergent selection occurred. Does it occur at contemporary or distant time periods? We therefore calculated the covariances of allelic frequency changes between the three forests for three different time periods (Figure 4):

- Contemporary time periods (Figure 4A)
- Adjacent time periods (Figure 4B)
- Distant time periods (Figure 4C)

**Figure 4.**
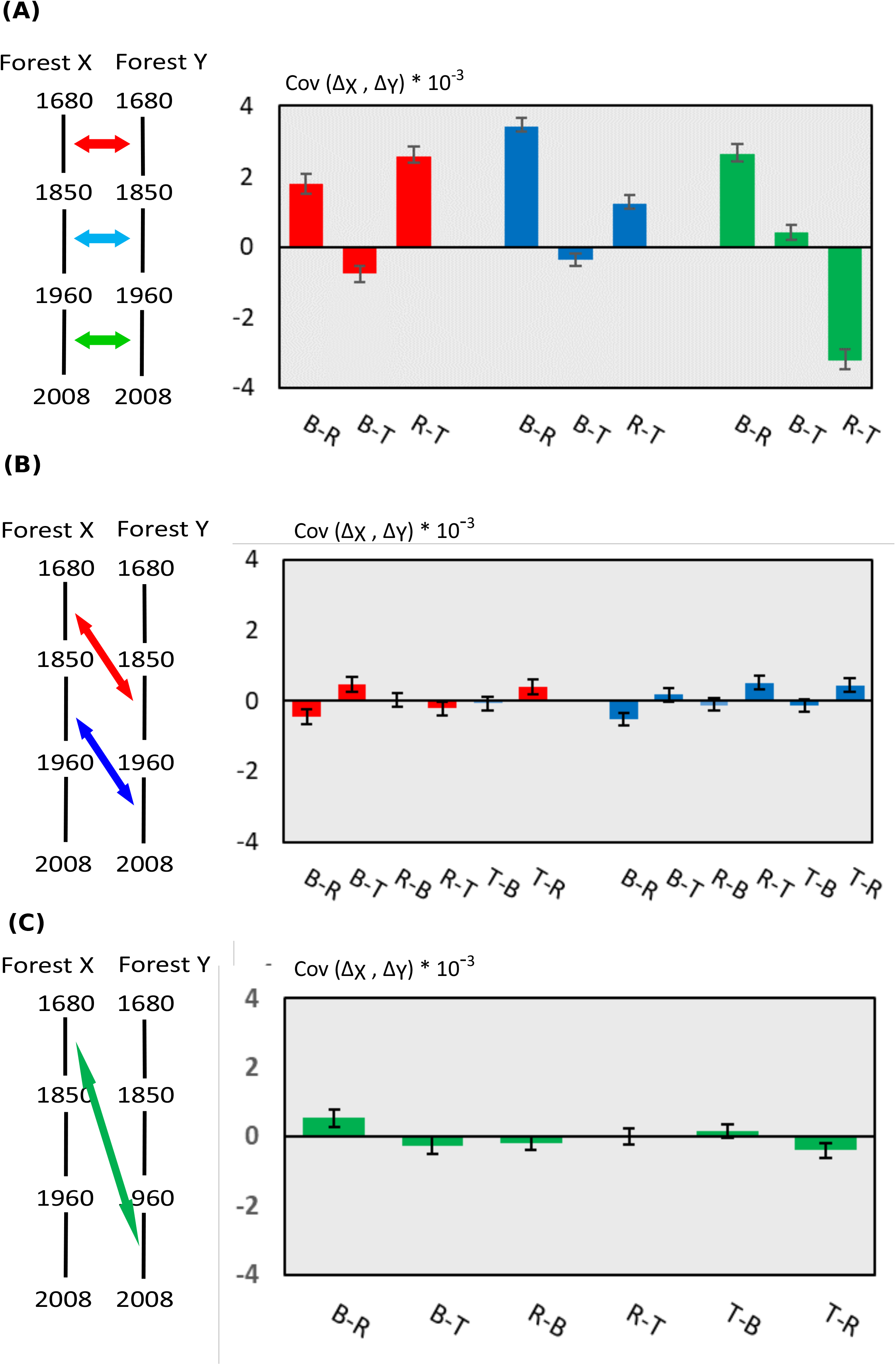
Temporal covariances of allelic frequency changes between the different forests for different time periods. Mean and 95% confidence intervals of the covariances were obtained by bootstrapping with 5000 iterations. Colours of the arrows on the left diagram indicate the time periods considered in the graphs. B : Bercé, R : Réno-Valdieu, T: Tronçais. (A) Temporal covariances of allelic frequency changes between forests for contemporary time periods. (B) Temporal covariances of allelic frequency changes between forests for adjacent time periods. (C) Temporal covariances of allelic frequency changes between forests for distant time periods.

We did not consider overlapping time periods, to avoid the inflation of covariances due to allelic frequency changes common to the covariance terms.

Overall, covariances between forests remained positive and significant for contemporary time periods, particularly between Bercé and Réno-Valdieu and between Réno-Valdieu and Tronçais (Figure 4A). Covariances between Bercé and Tronçais were close to zero regardless of the time period considered. Leaving these latter covariances aside, there was a trend over time towards a decrease in contemporary covariances from more ancient to more recent time periods (Figure 4A). The covariances between two forests, between adjacent or distant time periods, were strikingly different from contemporary covariances, as they were all close to zero (Figure 4B and 4C).

### Gene ontology and enrichment analysis

We divided all scaffolds into tiles of 100 kb in length and recalculated the temporal covariance of allele frequency changes for all tiles. We then identified putative regions (tiles) under linked selection on the basis of their covariances between the two earliest time intervals Cov(Δ_1680–1850_, Δ_1850–1960_) being above 0.01 in all three replicates. We restricted our analysis to the two earliest time intervals (yielding positive values in all three forests (Figure 3A), for which strong covariances were also observed between forests (Figure 4). In total we identified 104 tiles exhibiting temporal covariances above the threshold, which corresponded to 1% of the tiles (Figure S4 and S5). We inventoried 280 protein coding genes in these regions. Functional annotations of the 280 genes putatively under linked selection are summarized in Table S2. Briefly, out of these 280 genes, 248 received a GO annotation from *A. thaliana*, and two also received eggNOG GO terms, leaving 30 genes without a GO annotation.

The 280 genes revealed significant enrichments in the different gene ontologies (Table S3, S4, S5). Enrichment analysis identified several Biological Processes (BP), most of which are related to the “plant-type hypersensitive response”, “defense response to fungus”, “wax and cutin biosynthetic processes” and “anther dehiscence”, with higher connectivity between the two first terms which gather 15 and 13 genes respectively (Figure 5, Table S3). Fifteen genes belong to the group “defense response to fungus”, eight of which encode proteins similar to AT2G34930, a LRR disease resistance family protein, one was similar to AT3G59660 (BAGP1) required for fungal resistance, one was similar to AT5G64120 (PRX71) encoding a cell wall-binding peroxidase involved in lignification and important for fungal defense, two genes similar to AT3G51550 (FER), a plasma membrane-localized receptor-like kinase involved in fungal infection but also in flowering regulation, and three genes similar to AT1G02205 (CER1), which is associated with the production of stem epicuticular wax and pollen fertility. These multiple copies of *Arabidopsis* homologs correspond to tandem duplications located on chromosomes 3 (AT2G34930, 8 homologs), chromosome 12 (AT3G51550 (FER), 2 homologs) and on the unassigned scaffold Qrob_H2.3_Sc0000124 for AT1G02205 (CER1) (3 homologs), respectively (Table S2). Enrichment results highlight additional Biological Processes contributing to resistance to biotic and abiotic (Supporting Information 3). Furthermore the Molecular Function and Cellular Component ontology groups included as well genes involved in stress resistance (Supporting Information 3).

**Figure 5.**
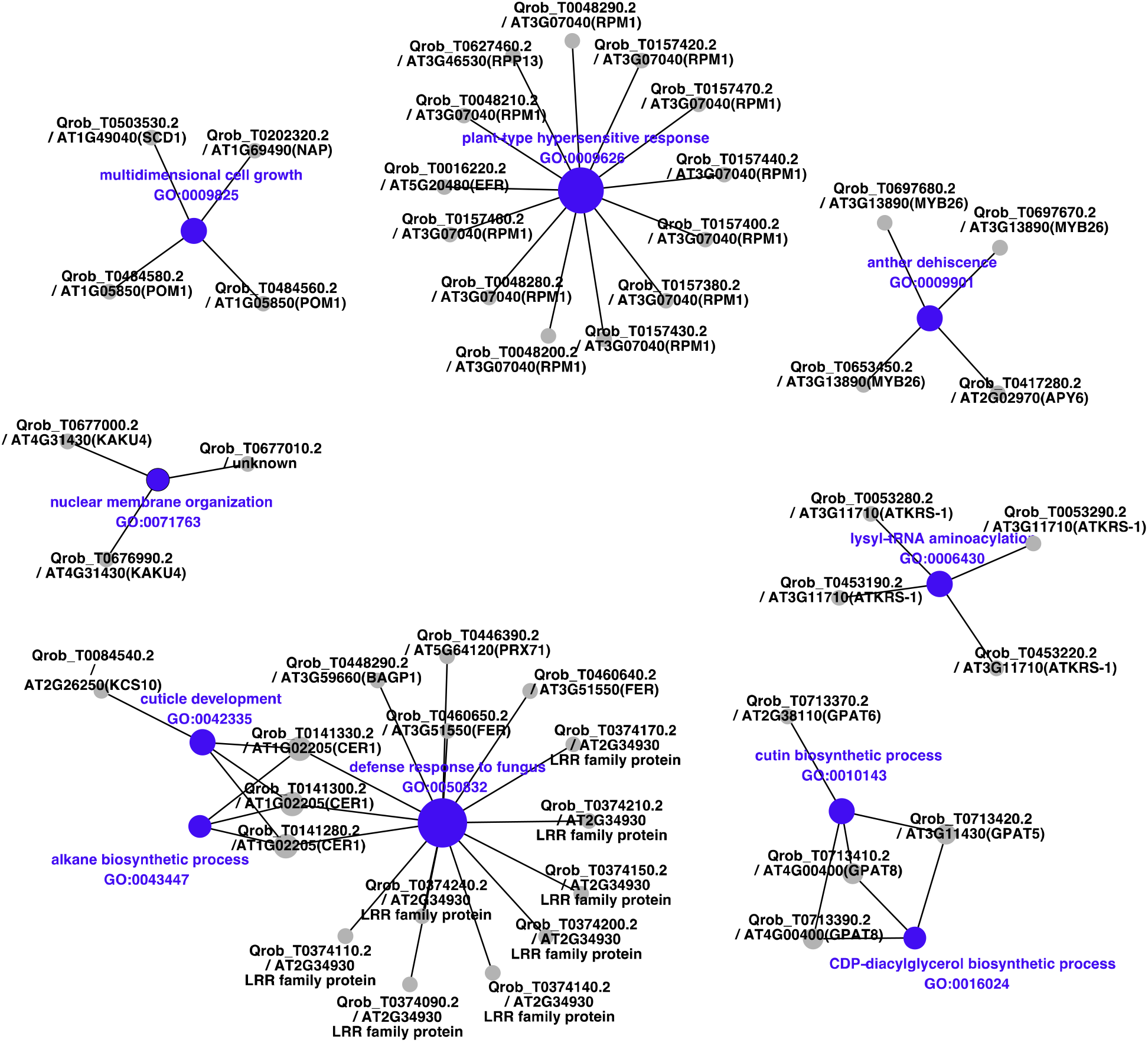
Network plot of the ten most significant Biological Process (BP terms from gene ontology enrichment analysis). The size of the nodes is proportional to their degree of connectivity. The labels correspond to the name of the *Quercus robur* gene model followed by the locus name of the best *Arabidopsis* homolog and the corresponding *Arabidopsis gene name* (from TAIR: https://www.arabidopsis.org/) between brackets when available. When no gene name exists, a short description (from TAIR or EggNOG databases) is added. If no description is available, the description is set to “unknown”.

## DISCUSSION

### Temporal covariances as imprints of genome-wide linked selection durin and after the LIA

Temporal allelic frequency changes for genes underlying adaptive polygenic traits have been studied in detail through theoretical approaches (Hollinger *et al*. 2019; Stephan & John 2020), simulations (Franssen *et al*. 2017), and empirical case studies (Long *et al*. 2015). Convergent predictions highlight small and subtle changes for genes targeted by directional selection, and have been supported by multi-generation selection experiments in model organisms (Burke *et al*. 2010; Barghi *et al*. 2019; Michalak *et al*. 2019). Our results suggest that genomic signatures of recent selection are also detected in natural populations of long-lived non-model species and over shorter time periods (or limited numbers of generations). Rapid evolution was reported at the genomic level in previous studies in non-model species in the context of rapid environmental change (Franks *et al*. 2018; Hamann *et al*. 2021) strong selection pressures (Dayan *et al*. 2019) or domestication (Guo *et al*. 2018), and this process was fueled by the presence of large amounts of standing genetic variation (Bitter *et al*. 2019; Dayan *et al*. 2019). The results we obtained in oaks meet theoretical expectations and show that tree populations are under genome-wide linked selection. Buffalo and Coop (Buffalo & Coop 2019) showed that, under directional selection (based on an exponential fitness function), the covariances of neutral allele frequency changes over time depend on the genetic variance of fitness (VA), recombination distance (R) and linkage disequilibrium, and found that the amount of covariances accumulated over a few generations was mainly determined by the compound factor VA/R. We recently reported that *Q. petraea* forest stands display substantial heritable differences in reproductive success (Alexandre *et al*. 2020). The genetic variance of fitness in a forest located between two of the forests studied here, Bercé and Réno-Valdieu, was 0.468 (Alexandre *et al*. 2020), which is at the upper end of the range of values reported in a recent review of the literature (89% of the reported values were below 0.20) (Hendry *et al*. 2018). In addition, most linkage disequilibrium is present in oak populations for physical distance lower that 5000 bp (Nocchi *et al*. 2021), which corresponds to 0.0052 cM taking into account the physical and genetic maps of oak (Bodenes *et al*. 2016; Plomion *et al*. 2018). The compound factor (VA/R) in oaks amounts therefore to 90, which should generate covariances varying between 0.002 and 0.004 after less than 10 generations (Buffalo and Coop (Buffalo & Coop 2019), their Figure 4, p.1019), which is within the range observed in our study (Figure 3A and 3B). Furthermore, QTL studies of fitness-related traits in oaks (growth, phenology and reproduction) have shown that all these traits depend on a large number of loci (Brendel *et al*.2008; Derory *et al*. 2010; Caignard *et al*. 2019), suggesting the genetic architecture of fitness is highly polygenic. Sessile oak and most likely trees in general are therefore prone to linked selection, provided that environmental constraints are strong enough for selection to operate.

Finally, the temporal allelic frequency changes assessed in our particular case study of age-structured cohorts are also potentially subject to other sources of variation that must be considered. Pollen flow between the age-structured cohorts within each forest may affect estimates of covariances. In the peculiar situation of age structured cohorts, gene flow between cohorts amplifies generation overlap. While the theory of temporal covariances generated by linked selection has been developed in the frame of discrete generation (Buffalo & Coop 2019), it is unknown how generation overlap may impact covariances. Given the age class distributions in even-aged stands, old cohorts occupy limited surface in comparison to mid age cohorts, and gene flow is likely more frequent within the latter then between different age classes. Furthermore gene flow between cohorts reduces temporal allelic changes between cohorts and would likely blur the covariances. Our covariances may, therefore, actually have been underestimated. Finally, the confounding effects of development-related changes in fitness may have contributed to increases in covariances too. If traits contribute to fitness changes over time in a long-lived species, then genetic covariances of fitness-related traits will increase between traits assessed at similar ages. This temporal serial autocorrelation — and its decay over time — have been reported for growth in pines (Kremer 1992). Such development-related covariances are entirely confounded with temporal covariances in our case. However, we would argue that developmental covariances occurring at the adult stage are likely to have a limited impact on temporal covariances, as selection is overwhelmingly more severe at the juvenile stage, with more than 90% of oak seedlings eliminated before the age of 10 years (Jarret 2004). Ultimately, human interferences are unlikely as well to have impacted temporal trends of the covariances, as these stands have been conducted under even-aged silviculture since the seventeenth century, and because selection is at its strongest during the juvenile period when no human mediated intervention takes place (Methods, paragraph “sampling”)

### Shifts of temporal covariances mirror climatic transitions between the LIA and modern times

If *Q. petraea* populations provide the necessary substrate for linked selection, what were the environmental constraints that triggered viability selection during the time periods considered? There is evidence that the LIA was a period with frequent extreme events in both summer and winter (Dobrovolny *et al*. 2010; Berchtold *et al*. 2012; Moreno-Chamarro *et al*. 2017) (Figure S2 and S3), these events being severe enough to affect natural populations of plants and animals. As illustrated in Supporting Information 2, extreme frosts and droughts were more frequent up to the late 19^th^ century than during more recent periods. Frost damage, including tree decline and deaths, was reported after extreme winters in the late LIA. (Hausendorff 1940) summarized historical writings about forest districts in Northern Germany mentioning recurrent episodes of adult oak tree mortality during the 1740-1748 period, following the extreme winter of 1739-1740 (Hausendorff 1940). Many historical documents from Catholic parishes reported extensive tree losses (fruit and forest trees, including walnuts, plum trees, oaks and chestnuts) following the winter of 1708-1709, which is known to be the most severe winter in European records (Avila & Avila 1987; Luterbacher *et al*. 2004). In Eastern France, a historical atlas dating from 1758 reported “The dead material was mainly oak. For example, forty six oak high forests belonging to all categories were totally dead.” This high mortality was presumably attributed to the extreme winter of 1708-1709 (Schnitzler 2020). However, the climate improved after 1850. There were only three reported extreme winters (1942, 1947 and 1963) and three extreme droughts (1921, 1934, 1976) during the 20^th^ century (Figure S2 and S3), during which oak decline was also recorded, but not the same extent as previously (Delatour 1983). These extreme events and the shift in their occurrence over time (during the second half of the 19^th^ century) may be responsible for generating the pattern of linked selection observed in this study.

As pointed out in the M&M part, selection intensity at the seedlings stage is extremely high and the size of the seedling population very large. Theory (Buffalo & Coop 2019) and empirical data from selection experiments (Buffalo & Coop 2020) suggest that, if the direction of selection is maintained over time, covariances tend to be positive, whereas shifts of directional selection are likely to decrease covariances, possibly resulting in negative values. In experimental selection, in which the direction of selection is maintained over successive generations, temporal covariances are predicted to decay as a result of recombination (Buffalo & Coop 2019). In natural populations, heritable fitness differences, the strength and shifts of direction of selection, and recombination are the drivers of the temporal covariances. The widespread and frequent occurrence of extreme frosts and drought events up to the late 19^th^ century in Europe may therefore be responsible for the positive covariance observed between the two earliest time periods (1680-1850 and 1850-1960, Figure 3A) within each forest and for the positive covariances at contemporary time scales between forests (Figure 4A). Conversely, the lower, and even negative covariances between the two more recent time periods (1850-1960 and 1960-2008, Figure 3B) may be a genomic signature of the change in the frequency of extreme events between the mid-19^th^ century and today. The maintenance of positive (or negative) covariances between adjacent time periods is reinforced by the limited impact of recombination in our study case. The short time interval between cohorts would have prevented opportunities for recombination events to break the linked polymorphisms. The time trends for covariances between forests support the interpretation that the strength and shifts of selection since the late LIA have shaped signatures of linked selection. Between-forest covariances were higher and more consistent across forests during the late LIA, when climatic conditions were harsher than the milder climatic conditions observed in the late 20^th^ century (Figure 4).

### Biotic interactions driven by adaptive response to climate

The functional analysis highlighted significant enrichments for genes involved mainly in plant defense responses to pathogens, or contributing to abiotic stress responses (temperature and drought) (Figure 5). Many of the genes encoding resistance proteins (R proteins) belong to the NBS-LRR families. These R-genes are widely represented in the oak genome, in which they appear as expanded groups (Plomion *et al*. 2018). R-proteins are involved in pathogen recognition and the subsequent activation of innate immune responses. Most R-proteins contain a central nucleotide-binding domain, the so-called “NB-ARC” domain, a TIR or CC N-terminal domain and a C-terminal LRR domain. The NB-ARC domain is a functional ATPase domain, and its nucleotide-binding state is thought to regulate the activity of the R protein. Such NBS-LRR receptors detect effectors used by pathogens to facilitate infection, and participate in the signaling cascade leading to various responses preventing further infection, such as the hypersensitive response, the production of reactive oxygen species and cell wall modification (Boyes *et al*. 1998; Roux 2010; El Kasmi *et al*. 2017). Interestingly two homologs of genes RPM1 and EFR were both involved in plant defense against *Pseudomonas syringae* (Boyes *et al*. 1998) which can infect a wide range of herbaceous and woody plants that have suffered frost and freezing damage (Young *et al*. 1988; Luisetti *et al*. 1991; Balestra *et al*. 2009). In addition, five genes encoding beta-glucosidases that could confer freezing/cold tolerance were identified (Thorlby *et al*. 2004; Fourrier *et al*. 2008; Ambroise *et al*. 2020). These genes may also participate in responses to other biotic/abiotic stresses (Baba *et al*. 2017; Vassao *et al*. 2018). Our functional analysis is consistent with the oak responses described in cases of oak decline ultimately leading to the death of the tree. Botanic and pathological descriptions of oak decline during the LIA are lacking, but reports of recurrent sparse oak dieback in more recent decades can be used to retrace the steps leading to oak death following severe winter or drought events. In a review of oak decline in Europe, Thomas et al. (2002) highlighted the combined effects of climatic extremes (drought or frost) and defoliating insects and pathogenic fungi (Thomas *et al*. 2002). The starting point is an extreme climatic event that weakens the trees (Vanoni *et al*.2016), which is then followed by pest and insect attacks, which ultimately kill the tree. For example, episodes of oak decline in the first half of the 19^th^ century in various parts of Europe were caused by a combination of winter frost, summer drought, insect defoliation (caused by oak leaf roller (*Tortrix viridana*) and oak processionary moth *Thaumetopoea processionea*),and several pathogenic fungi (mainly powdery mildew *Erisyphe alphitoides*) and root pathogens (*Armillaria* and *Phythopthora*) (Delatour 1983; Donaubauer 1998; Thomas *et al*.2002). The effect of drought or frost driving pathogen and insects dynamics has been reported in other oak species, e.g. in Mediterranean oaks (Colangelo *et al*. 2018) and American red oaks (Wood *et al*. 2018). In summary, extreme events, such as severe frost or drought, expose trees not only to abiotic stresses, but also to biotic selection pressures. These, in turn, trigger resistance responses, which were identified at genomic level in the functional analysis. We suspect that these processes also operated during the late LIA and that exposure to selection was more stringent, leading more widespread death (Hausendorff 1940; Schnitzler 2020), and, ultimately, to convergent linked selection across the three forests.

## CONCLUSION

Our results clearly show that selection has been operating at recent and contemporary time scales in long-lived species such as oaks, but in different directions during cold and warm periods. Patterns of covariance changes indicate that selection intensity and direction fluctuated during the last three centuries, reflecting changes in climatic conditions. Evolution over short time periods may play a more important role in trees than previously thought, but its effects may be partially erased over longer periods if selection fluctuates. Beyond oaks, other woody species sharing similar life history traits and attributes (large standing genetic, polygenicity of fitness, high fecundity and severe selection screening at the juvenile stage..) may as well be prone to linked selection and allow retrospective tracking of evolutionary and adaptive pathways. Such retrospective approaches may improve our understanding of future responses to ongoing climatic changes. Finally our findings may lead also to practical silvicultural implications. Our conclusions raise concerns about the maladaptation of old centennial trees under current climate conditions. They were exposed to different selection pressures driving different adaptive responses. And today they may swamp younger oak stands by maintaining a temporal ”migration load” by pollen flow. Our findings argue therefore for a decrease of generation time in managed oak forest to overcome maladaptation of old extant cohorts and “migration load”.

## Acknowledgements

This research was supported by the European Research Council through an Advanced Grant (project TREEPEACE # FP7-339728), by France Génomique (project EVOL-OAK, ANR-10-INBS-09–08), and by the French Forest Service (ONF) (INRAE-ONF TREEPEACE contract). Jun Chen was financed by the National Natural Science Foundation of China (31972946). We thank the staff of the ONF Research and Development Department (Myriam Legay, Lucie Arnaudet) and the ONF staff at the National Forests of Bercé, Réno-Valdieu and Tronçais (Anthony Jeanneau, Vincent Breton, Benjamin Laurendeau, Loïc Nicolas, Alexandre Durin) for technical assistance during the sampling of cohorts and trees. We acknowledge the contribution of Niklaus Zimmermann (WSL, Birmensdorf, Switzerland) for downscaling the historical climatic data. We thank Leo Arnoux, Sylvain Delzon, Patrick Reynet, and Florine Routier (UMR BIOGECO) for collecting bud, leaf and cambium samples for the 12 cohorts. We thank Alexis Ducousso (UMR BIOGECO) for assistance with forest selection, and Benjamin Brachi and Santiago Gonzalez-Martinez (UMR BIOGECO) for fruitful discussions during data analysis. We thank the Genotoul Bioinformatics Platform (Bioinfo Genotoul) at Toulouse (France) for providing computing and storage resources, and the Genoscope (CEA, Commissariat à l’Energie Atomique et aux Energies Alternatives) at Evry (France) for sequencing the oak samples. We thank Vince Buffalo for his helpful comments on an earlier version of the manuscript.

## Authors contribution

Conception and coordination of the study: A.K., C.P., M.L.; Sampling of forests and cohorts: A.K., F.M., L.T.; Collection of samples and DNA extraction: L.T., B.D., C.L., A.K., T.L., D.B.; Dendrochronological analysis: D.B., F.L.; Whole genome sequencing: J.M.A., K.L., C.P.; Bioinformatic analysis: D.S., I.L., J.C., T.L.; Enrichment analysis: J.C.L.; Data analysis: D.S., J.C., A.K., M.L.; Writing of the manuscript: D.S., J.C., J.C.L., A.K., M.L., T.L. All authors reviewed the manuscript.

