## Supporting Information for "Genome-wide evolutionary response of European oaks since the Little Ice Age"

**Supporting Information S1**

Even-aged oak forests and age structured cohorts

**Supporting Information S2**

Climatic trends over the studied period in the three forests.

**Supporting Information S3**

Enrichment analysis of genes located in outlier tiles.

**Supporting Information S1**

**Even-aged oak forests and age structured cohorts**

Most sessile oak stands in France are managed under even-aged regimes (Jarret 2004). Under such regimes, a forest is subdivided in a number of parcels (compartments) comprising trees that are approximately the same age, and all age classes are evenly distributed in different compartments. We sampled three forests located in the western part of France (Tronçais, Bercé and Réno-Valdieu (Figure 1A), where the rotation age is about 200 years and thus comprising compartments of different ages from 0 to 200 years. However within each of these forests, foresters have also maintained "older" age class compartments for patrimonial and historical reasons. In particular compartments contemporary of the first implementation of even-aged regimes that took place in the mid 17$^{th}$ century by the statesman Colbert were maintained (Gallon 1752). We sampled 4 compartments in each forest that were approximatively 340, 170, 60 and 12 years old (born approximatively in 1680, 1850, 1960 and 2008), which we will later call age structured cohorts (Table S1, Figure 1B). The area of a compartment varies between 10 to 30 ha (Figure1B) and thus comprises census sizes of about 2000 to 6000 trees for the oldest cohorts. These numbers increase for younger cohorts and may reach up $1*10^6$ to $3*10^6$ in the youngest stages. On average, compartments corresponding to the different cohorts were separated by less than 2 kms (Figure 1A) and were also located at similar elevations (Table S1), thus limiting microenvironmental differentiation between cohorts, and maintaining potential gene flow among cohorts. All sampled trees were georeferenced, and an additional limited number of trees were felled for age assessments (Figure 1B). Dendrochronological age assessments matched with management documents. The period of establishment of a cohort was about 30 years for the oldest cohorts, which corresponds to the time lag that was necessary for renewing a stand by natural regeneration. In more recent times regenerations durations extended over 10 to 20 years (Figure 1B).

| Forest | Cohort | Compartment# | Latitude | Longitude | Variation elevation (m) |
|---|---|---|---|---|---|
| **Bercé** | B1 | 227 | 47.78582 | 0.48701 | 161 to 173 |
| (5405ha) | B2 | 157 | 47.80993 | 0.39948 | 171 to 179 |
| Lat :47.8092 | B3 | 255 | 47.77766 | 0.49389 | 151 to 161 |
| Long :0.4004 | B4 | 226 | 47.78672 | 0.49221 | 155 to 171 |
| **Réno-Valdieu** | R1 | 39 | 48.52466 | 0.67761 | 243 to 258 |
| (1645ha) | R2 | 38 | 48.52760 | 0.67888 | 240 to 245 |
| Lat :48.5146 | R3 | 59 | 48.51511 | 0.67110 | 246 to 264 |
| Long: 0.6706 | R4 | 44-45 | 48.52579 | 0.66189 | 243 to 256 |
| **Tronçais** | T1 | 152 | 46.67122 | 2.77253 | 243 to 274 |
| (10600ha) | T2 | 62 | 46.65991 | 2.80044 | 265 to 272 |
| Lat :46.6598 | T3 | 150 | 46.67754 | 2.78378 | 252 to 267 |
| Long:2.7059 | T4 | 234 | 46.65815 | 2.70625 | 227 to 267 |

**Table S1.** Geographic coordinates of sampled forests and age structured cohorts

**Supporting Information S2**

**Climatic trends over the studied period in the three forests.**

Temperature reconstructions of the last millennium indicate substantial changes (Corona *et al.* 2010; Luterbacher *et al.* 2016; Anchukaitis *et al.* 2017; Neukom *et al.* 2019; Wang *et al.* 2019). A consistent trend towards warmer climates during the Medieval period, followed by a much cooler period between approximately 1450 and 1850 has been reported. This 400 year-long period is described as the "Little Ice Age" (LIA) (Matthes 1939; Tkachuck 1983). Mean summer temperatures decreased by 0.5°C to 2°C between the Medieval Warm Period and the Little Ice Age, and then increased again, by more than 1.5°C, to reach current values (Corona *et al.* 2010; Anchukaitis *et al.* 2017). The causes of the LIA remain a matter of debate (Nesje & Dahl 2003; Crowley *et al.* 2008; Palastanga *et al.* 2011; van Oldenborgh *et al.* 2013; Ruddiman *et al.* 2016; Owens *et al.* 2017), but the impact of this period on agriculture and society at large is clear (Pfister 1984; Fagan 2002; Parker 2013).

We used regional and temporal high-resolution (1 km coarse resolution) temperature reconstructions based on a combination of instrumental data, documentary records and ice core and tree ring proxy data by (Luterbacher *et al.* 2004) to monitor retrospectively temperature changes in the three forests since the mid seventeen century (Figure S1). These data illustrate the temperature trends that have been reported earlier since the LIA. The late LIA was dominated by two harsher periods which occurred during the late seventeen century overlapping with the Maunder Minimum (MM) solar activity (1645-1715, (Owens *et al.* 2017)) and during the mid-nineteenth century (1840 to 1860) a period that was preceded by intense volcanic activity was reported (Crowley *et al.* 2008; Bronnimann *et al.* 2019) (Figure S1).

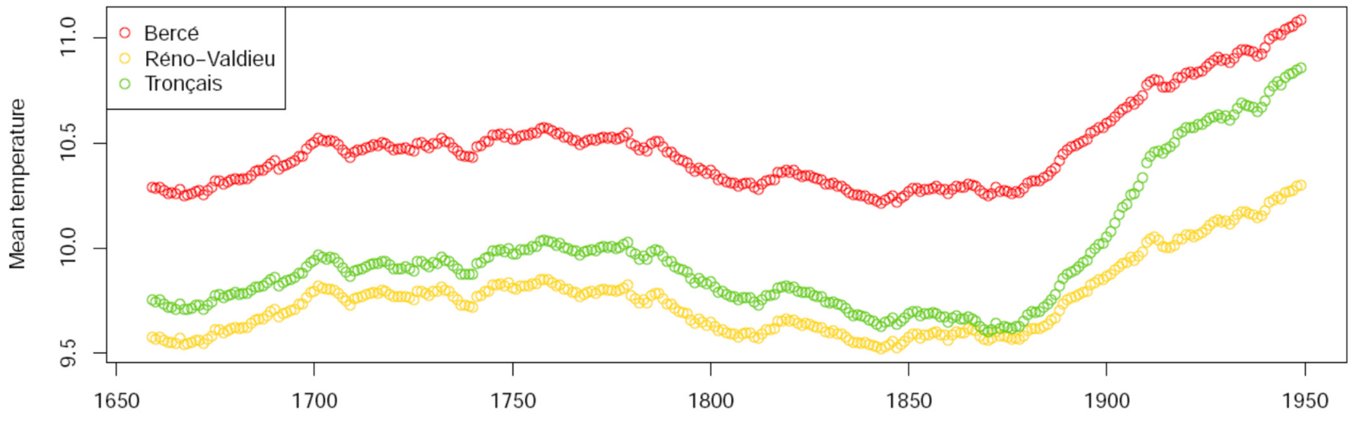


**Figure S1**. Mean yearly temperature trends in the three studied forests.
Temperatures were reconstructed according to Luterbacher (Luterbacher et al., 2004) using a combination of instrumental data, documentary records and ice core and tree ring proxy data. Yearly means were smoothed over 50 years.

There is also observational evidence of an increased frequency of extreme cold winters during these two periods based on instrumental temperatures recorded at the Observatory of Paris between 1676 and 2010 (Rousseau 2012) (Figure S2, Figure 3D). Occurrences of extreme winters during that period are also supported by historical documents. In his seminal work Le Roy Ladurie mentions that 1690 -1700 was the coldest decade in modern history with extreme winters occurring in 1695 to 1698 ((Le Roy Ladurie 2004), p.437, 487) and 1708-1709 is the worst winter ever reported ( (Le Roy Ladurie 2004) p.514 therein; (Luterbacher *et al.* 2004)). Similarly the decade 1840-1850 is a period where 3 extreme winters (1841, 1845, 1847 in addition to 1830 and 1838) were reported in Rousseau's survey (Rousseau 2012).

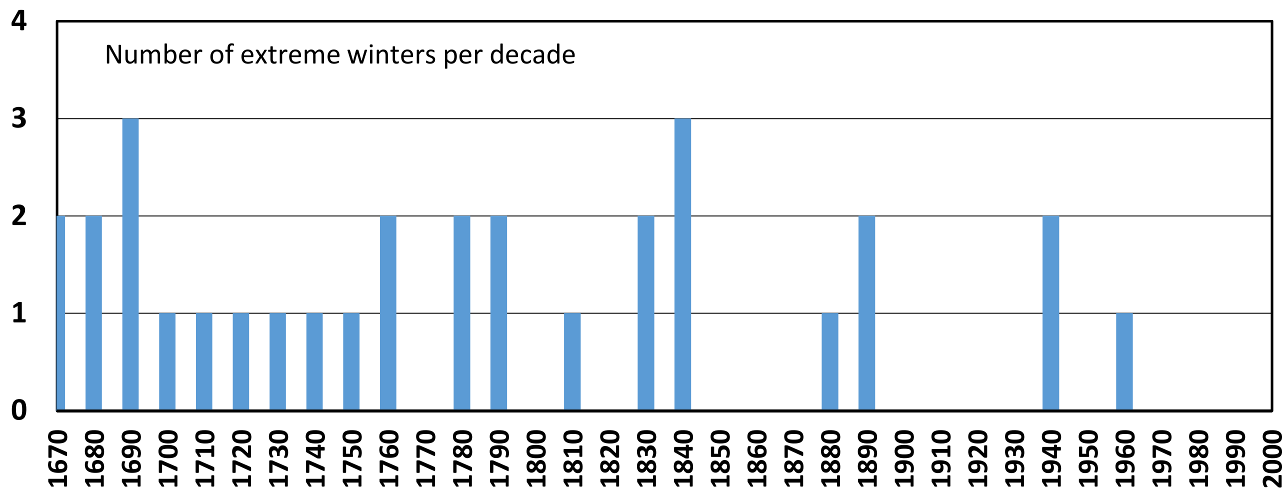


**Figure S2.** Number of extreme winters per decade.

Data compiled from Instrumental temperatures recorded at the Observatory of Paris between 1676 and 2010 (Rousseau, 2012). Winters were considered as extremes when the mean winter temperature (December-January-February) was lower by more than 2°C to the mean winter temperature during the 19$^{th}$ century in Paris.

Similarly to extreme winters we examined also occurrences of extreme summer droughts by mining Cook's data base of Old World megadroughts (Cook *et al.* 2015). This data base provides hydroclimatic reconstructions based on instrumental, historical, archaeological, and natural records during the last millennium. The data base allows to draw year to year regional maps of Palmer Drought severity indices (Palmer 1965; van der Schrier *et al.* 2013) (Figure S3, Figure 3D). Occurrences of extreme summer droughts mirror to some extent also occurrences of extreme winters, thus confirming that the LIA was not just a cooler period but also an age of climatic extremes (Fagan 2002; Le Roy Ladurie 2004, 2006; Blom 2019). While there was a clear overlap of extreme summers and winters during the two most severe periods of the LIA (late seventeenth century and mid-nineteenth century, Figure 3D), the overall frequency of extreme events decreased notably after 1850. The two oldest age structured cohorts are almost synchronous to the two most severe periods recorded during the LIA. Cohort 4 in the three forest originated during the late seventeenth century while trees of cohort 3 established in the mid-nineteenth century (Figure 3D, Figure S2 and S3).

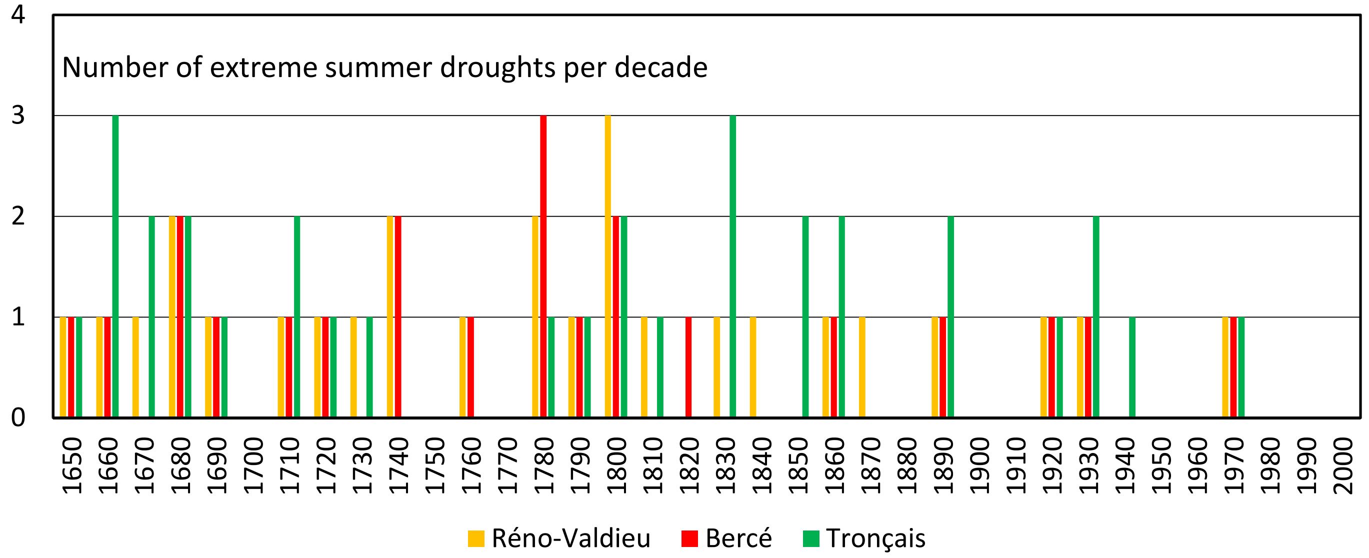


**Figure S3**. Number of extreme summer droughts per decade near each studied forest according to Cook's data base of Old World megadroughts (Cook et al., 2015).

Summers were considered as extremes when the Palmer's drought severity index (PDSI), which is a measure of soil moisture availability, was lower than -4 (Palmer, 1965; Van der Schrier et al., 2013). According to Palmer's classification of drought based on PDSI, values lower than -4 correspond to "extremely dry" conditions, which is also the most extreme class of PDSI.

**Supporting Information 3**

**Enrichment analysis of genes located in outlier tiles.**

In addition to the two first Biological Processes described in the text, two other groups gather genes potentially involved in resistance to biotic or abiotic constraints (Figure 5, Table S3): “Cutin biosynthetic process” and “anther dehiscence”. “Cutin biosynthetic process” included four genes: two genes similar to AT4G00400 (GPAT8) and two other similar to AT2G38110 (GPAT6). Cutin is a polymer that is partly covered and interspersed with waxes to form the cuticle, a physical barrier that protects the plant against water loss, irradiation, xenobiotics, and pathogens (Serrano *et al.* 2014).

A total of four genes were associated with the “Anther dehiscence” GO term: one was similar to AT2G02970 (APY6) involved in pollen exine pattern formation and anther dehiscence, and three similar to AT3G13890 (MYB26), which is known to control cellulosic secondary wall thickening in the endothecium. Only two of the three copies were tandem duplicates.

The term “ADP binding” was the most significant Molecular Function ontology group recognized in the enrichment analysis (Table S4). Most of the genes in this category encode proteins carrying NB-ARC or LRR domains and one transmembrane receptor, including the 12 oak homologs of RPM1 and RPP13 described above. This finding is consistent with the importance of the nucleotide binding site (NBS) and ATP/ADP binding in pathogen sensing(DeYoung & Innes 2006). Finally, “extrinsic component of plasma membrane” is the most significant Cellular Component GO terms (Table S5). It gathers the 12 oak homologs of RPM1, consistent with previous results.

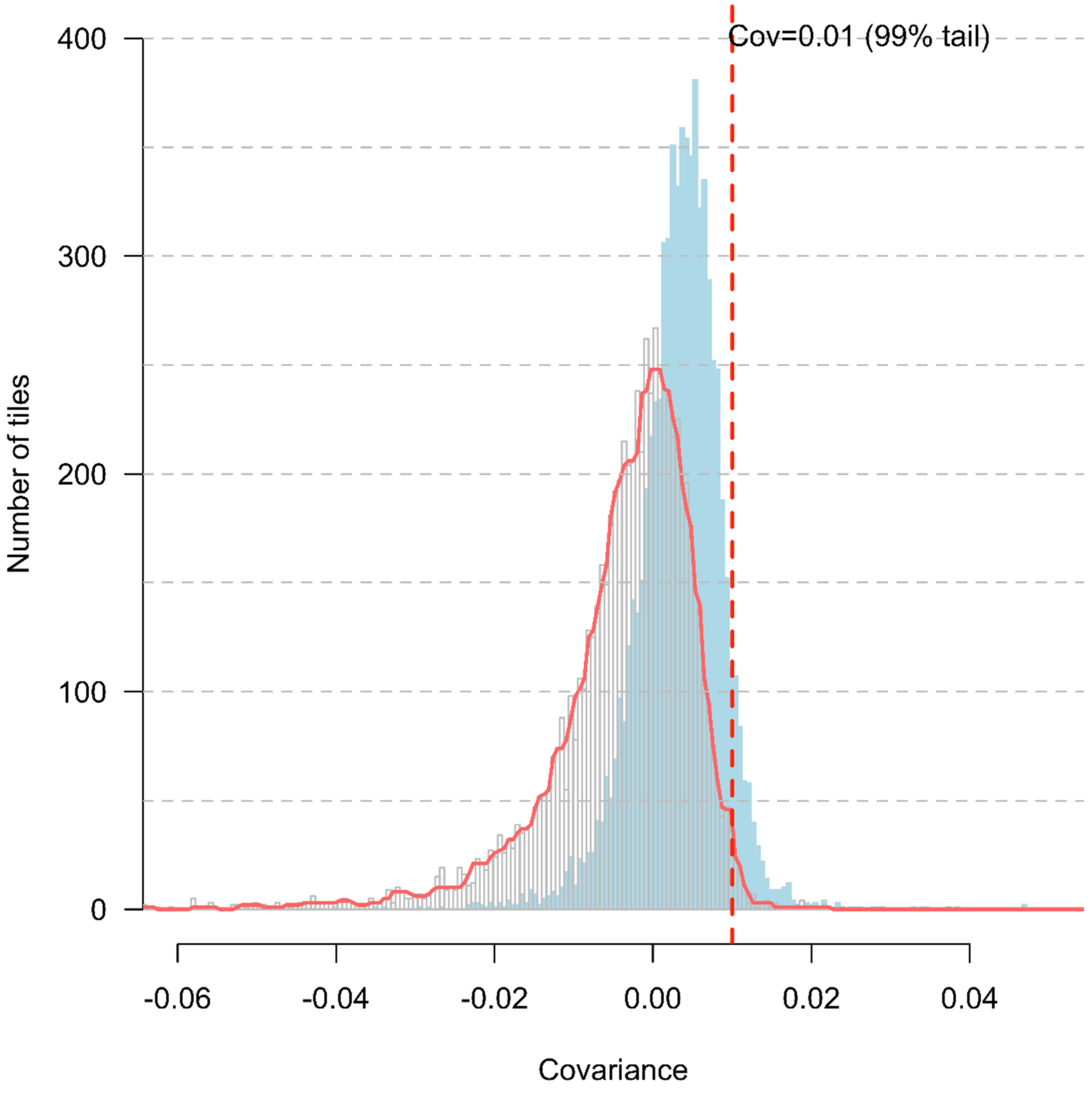


**Figure S4.** Distribution of the temporal covariances between allelic frequency changes between the two oldest time periods (*Cov*($\Delta_{1680\text{-}1850}$, $\Delta_{1850\text{-}1960}$)), calculated for tiles of 100kb.

The distribution corresponds to the covariances calculated at the tile level. The red distribution corresponds to the minimum values between the three forests ; the red dotted cut off line (0.01) is the threshold used to select the outliers tiles (104 tiles which represent 1% of the tiles). The blue shaded distribution corresponds to the mean covariances across the three forests.

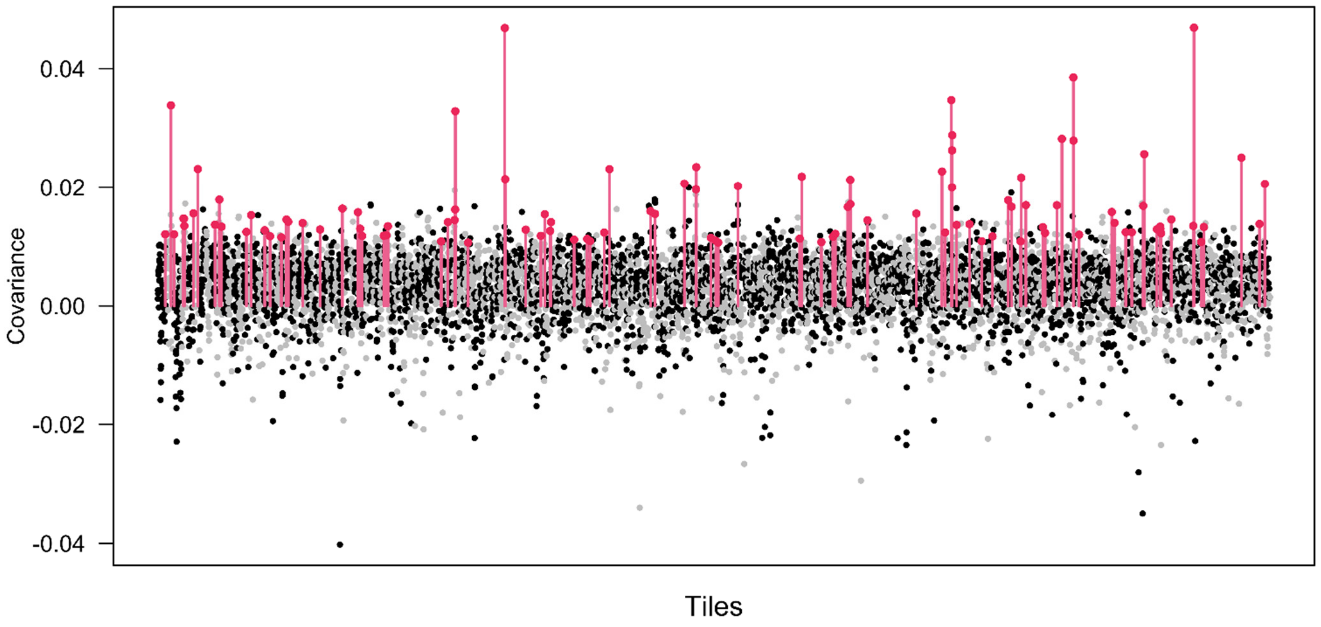


**Figure S5.** Manhattan plot of the temporal covariances between allelic frequency changes between the two earliest time periods ($Cov(\Delta_{1680\text{-}1850}, \Delta_{1850\text{-}1960})$) calculated at the tile level, over the whole genome.

Data correspond to the tile mean covariances across the three forests. Grey and black data points correspond to different consecutive scaffolds. Red lines correspond to the outlier tiles.
